# Cannabinoid Receptor 2 Activating Antibodies: A Promising Therapeutic Strategy for Macrophage-Driven Fibro-Inflammatory Diseases

**DOI:** 10.1101/2025.07.11.664477

**Authors:** Raghavender Reddy Gopireddy, Dipankar Bhattacharya, Swastik Sen, Monica A. Schwartz, Scott L. Friedman, Richard C. Yu, Toshi Takeuchi, Lauren J. Schwimmer

## Abstract

Fibrotic diseases, often fueled by chronic inflammation, represent a major unmet medical need. A critical driver of this process is the activation of macrophages, which secrete pro-inflammatory cytokines and chemokines, leading to immune cell recruitment and collagen deposition by myofibroblasts. Agonizing the cannabinoid receptor 2 (CB2), expressed on macrophages, offers a potential therapeutic avenue to suppress this activation. Despite widespread interest in CB2 modulation, challenges remain in the development of small molecule agonists, including insufficient specificity and poor drug-like properties. Antibodies are highly specific with favorable pharmacokinetics and bioavailability, but GPCR agonist antibodies have been difficult to discover. Here we present first-in-class CB2 agonist antibodies, AB120 and AB150. These antibodies are CB2 specific, G-alpha biased, and effectively decrease the expression of macrophage activation markers and key pro-inflammatory cytokines, including IL-6, IL-1β and TNF-α. Furthermore, using an ex vivo human precision-cut liver slice (hPCLS) model of liver fibrosis, we observe that treatment with these CB2 agonist antibodies not only reduces inflammatory markers but also decreases collagen expression, indicating a potential to halt or reverse fibrosis. These novel CB2-specific agonist antibodies hold promise as therapeutics for a range of fibrotic and inflammatory conditions driven by chronic macrophage activation and demonstrate the potential of GPCR antibody agonists to drug this challenging class of targets.

**Significance Statement:** Fibrotic diseases driven by chronic inflammation lack effective treatments. CB2 agonism modulates macrophage activation to reduce pro-inflammatory cytokines suggesting a novel avenue of treatment, but small molecule CB2 agonists have faced substantial limitations due to off-target effects. We introduce two novel CB2-specific agonist antibodies with potent anti-inflammatory and anti-fibrotic effects in vitro and in a human ex vivo liver fibrosis model. These antibodies provide a promising new therapeutic strategy and highlight the advantages of antibodies as GPCR agonist drugs.

## Introduction

The cannabinoid receptor 2 (CB2) has received notable therapeutic interest for a range of conditions, particularly those involving inflammation and immune modulation.^1–5^ However, small molecule agonist development has been hindered by the high lipophilicity of the molecules, resulting in poor pharmacokinetics, and by cross-reactivity to CB1, which is highly expressed in the central nervous system (CNS) and associated with adverse psychoactive effects.^6–8^ Antibodies are stable, well-expressed, have excellent pharmacokinetic properties, and have exquisite specificity toward their targets.^9^ Therefore, a CB2 antibody agonist is poised to overcome the major challenges encountered by small molecule agonists. Here, we functionally characterize the first reported CB2 antibody agonists, AB120 and AB150,^10,11^ which demonstrate selective CB2 activation and anti-inflammatory properties.

CB2 is a class A G-protein coupled receptor (GPCR) that is primarily expressed in lymphocytes and myeloid cells, including macrophages, microglia, and peripheral sensory neurons.^12,13^ CB2 couples to G_i/o_ and modulates various intracellular signal transduction pathways, including inhibition of cAMP production, ERK1/2 phosphorylation and β-arrestin-2 recruitment.^8,14,15^ These signals ultimately modulate expression of anti-inflammatory and anti-fibrotic genes.^16–22^ CB2 activation induces macrophage autophagy and suppresses macrophage activation,^23–25^ thereby reducing expression of pro-inflammatory cytokines and chemokines including IL6, TNF-α, IL-1β and CCL4. Chronic macrophage activation can lead to fibrosis in many tissues, including the liver.^22,26^ Therefore, CB2 agonists are well situated to treat chronic inflammation and fibrotic diseases in which macrophages contribute to the pathology.

Selective small molecule CB2 agonists have been pursued for treating various conditions such as inflammatory pain, neuropathy, and liver fibrosis.^4,5,8,22,27^ While these compounds primarily target CB2, their development has been hindered by the off-target effects on CB1,^16,28^ as the high conservation of the ligand binding pocket makes it challenging to produce highly specific small molecules.^29^ Specificity for CB2 is crucial, as even low CB1 receptor occupancy (4%-14%) can lead to activation of the receptor and subsequent inhibition of locomotor activity in mice.^30^ Off-target activation of CB1 is undesirable as CB1 is widely expressed in the CNS, where activation leads to the psychoactive effects associated with the recreational use of cannabinoids including increased appetite.^31^ In addition, activation of CB1 can be pro-fibrotic, countering the biological effect of the CB2 agonism.^32,33^

Due to off-target liabilities of some small molecule drugs, highly specific and selective antibody drugs are attractive alternatives and are now an important component of the therapeutic landscape.^34^ The CB2 orthosteric binding pocket is highly conserved, while the low amino acid sequence conservation of the surface available residues between CB1 and CB2 (26% identity) makes it unlikely that CB2 agonist antibodies would cross-react to CB1. Thus, antibodies that target the extracellular loops of CB2 and activate the receptor allosterically have the potential to be more specific than small molecules.

Here we demonstrate in vitro cAMP, β-arrestin and ERK1/2 signaling of our CB2 agonist antibodies, AB120 and AB150, in mouse and human macrophage cell lines. This activity is specific to the CB2 receptor and biased toward cAMP over β-arrestin and ERK1/2. We also show that these CB2 agonist antibodies reduce macrophage activation and decrease secretion of pro-inflammatory and pro-fibrotic proteins from cell lines and precision cut human liver slices (hPCLS). AB120 and AB150 have potential therapeutic applications in a variety of inflammatory conditions where macrophage biology is implicated, including peripheral neuropathies and fibrotic diseases.

## Results

We performed in vitro assays with the CB2 agonist VHH-Fc chimeric antibodies AB120 and AB150 using mouse (RAW264.7) and human (THP-1 differentiated with PMA, Sup Fig 1) macrophage cell lines, which express both CB1 and CB2^35^ to determine the functional activity and selectivity for CB2 and not CB1. The cannabinoid receptors are primarily G_i_ coupled, whereby activation leads to inhibition of adenylate cyclase and reduction of cAMP production. We utilized a bioluminescence-based high-throughput assay to measure cAMP, in which the cell lines were stimulated with NKH477 (a forskolin derivative) to produce cAMP and treated with AB120, AB150, HU-308 (CB2 small molecule agonist),^8,27^ or AB100 (negative isotype control antibody). Importantly, both, AB120 and AB150, but not AB100, potently inhibit cAMP production in both mouse and human macrophage cell lines. The EC_50_ and E_max_ of both antibodies are comparable to HU-308, which was chosen as the CB2 agonist benchmark due to its >100-fold CB2 selectivity over CB1 and balanced activity across multiple signaling pathways.^8^ (Figure 1A and D, Table 1). This agonist activity is inhibited by SR144528,^36,37^ a CB2 selective antagonist (Figure 1B and E), but not by SR141716A,^38^ a CB1 selective antagonist (Figure 1C and F), which can inhibit ACEA, a CB1 selective agonist. These data demonstrate that the effect on cellular cAMP levels by AB120 and AB150 is specific to CB2 receptor agonism.

**Fig.1.**
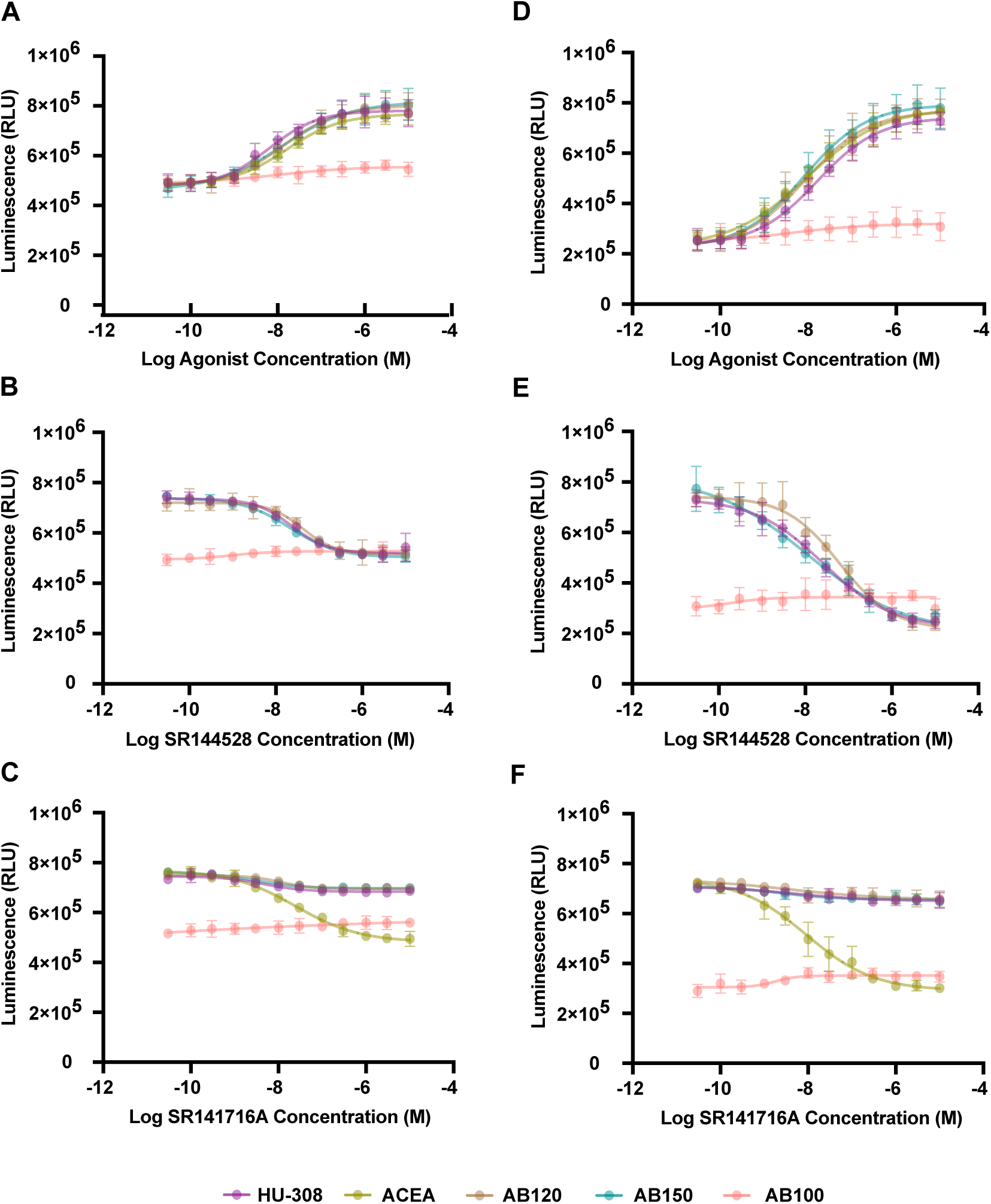
Dose response curve for CB2 specific agonist-dependent cAMP inhibition in RAW26.7 (A-C) cells and THP1-cells (D-F). CB2 specific agonist (HU-308), CB1 specific agonist (ACEA), and CB2 specific agonist antibodies (AB120, AB150) decreases NKH477 induced cAMP production as detected via cAMP-Glo luminescence (Promega), where cAMP concentration is inversely proportional to luminescence (A and D). Additionally, CB2 selective antagonist, SR144528, inhibits HU308, AB120 and AB150 agonist activity (B and E). CB1 selective antagonist, SR141716A, only inhibits CB1 selective agonist, ACEA, activity (C and F). Data are represented as mean ± SD from two independent experiments.

**Table 1.**
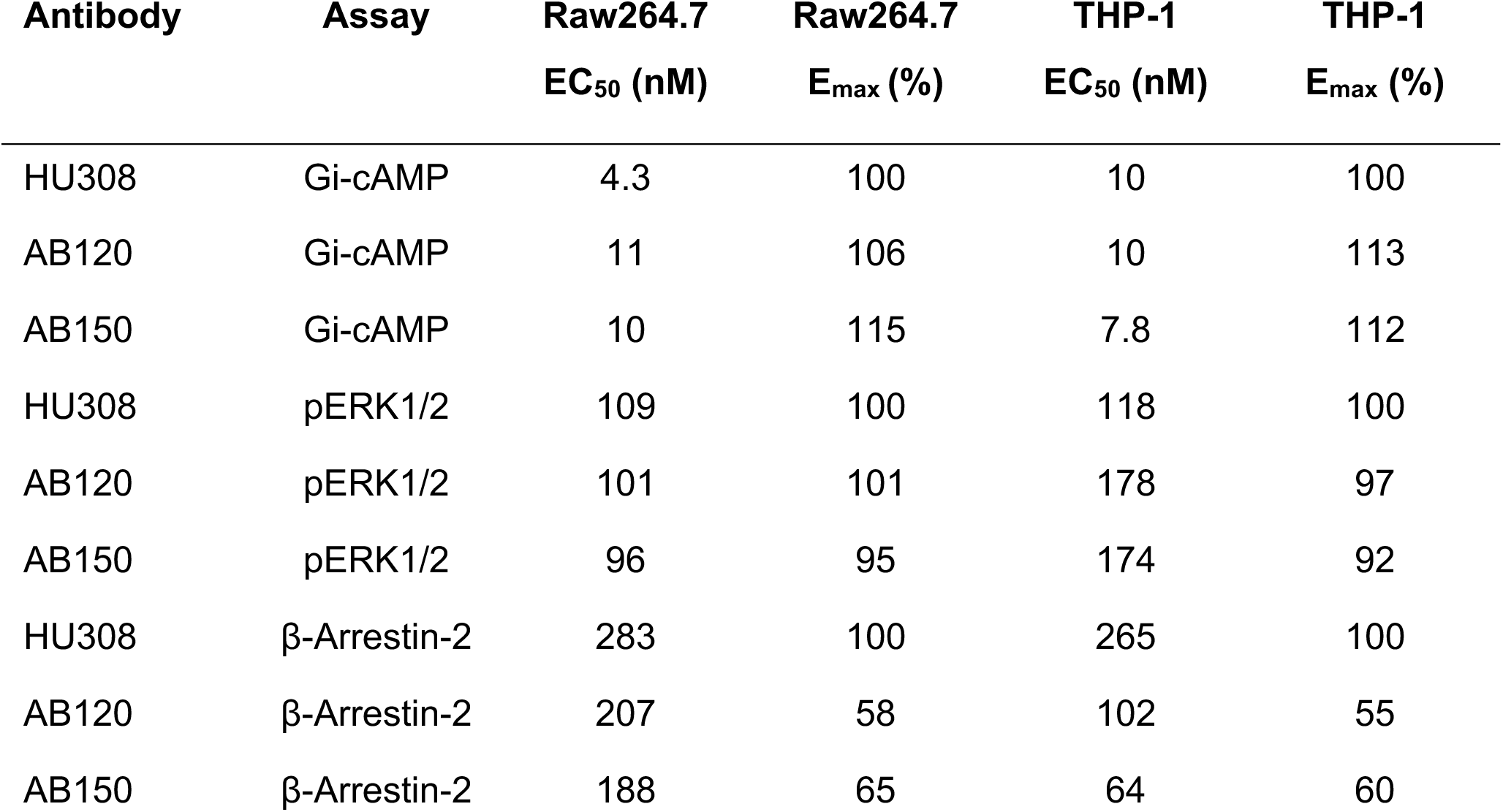
Efficacy and Potency profile of CB2 agonists.

CB2 agonism may also induce recruitment of β-arrestin-2, leading to internalization and downregulation of CB2,^39^ potentially limiting the in vivo effect of the antibodies. Therefore, it is important to characterize this signaling pathway. A TR-FRET based high-throughput assay was used to assess β-arrestin-2 recruitment by measuring the interaction between β-arrestin-2 and AP2, which interacts with β-arrestin-2 only after engagement with CB2. We found that, although AB120 and AB150 have similar potencies to HU-308 for β-arrestin-2 recruitment on both human and mouse macrophages (Figure 2 and Table 1), both are partial agonists with lower E_max_ compared to HU-308 in this pathway. This demonstrates that both antibodies are also biased agonists, as the potencies (EC_50_) for the G-protein dependent cAMP inhibition are >10-fold more than that for β-arrestin-2 recruitment.

**Fig. 2.**
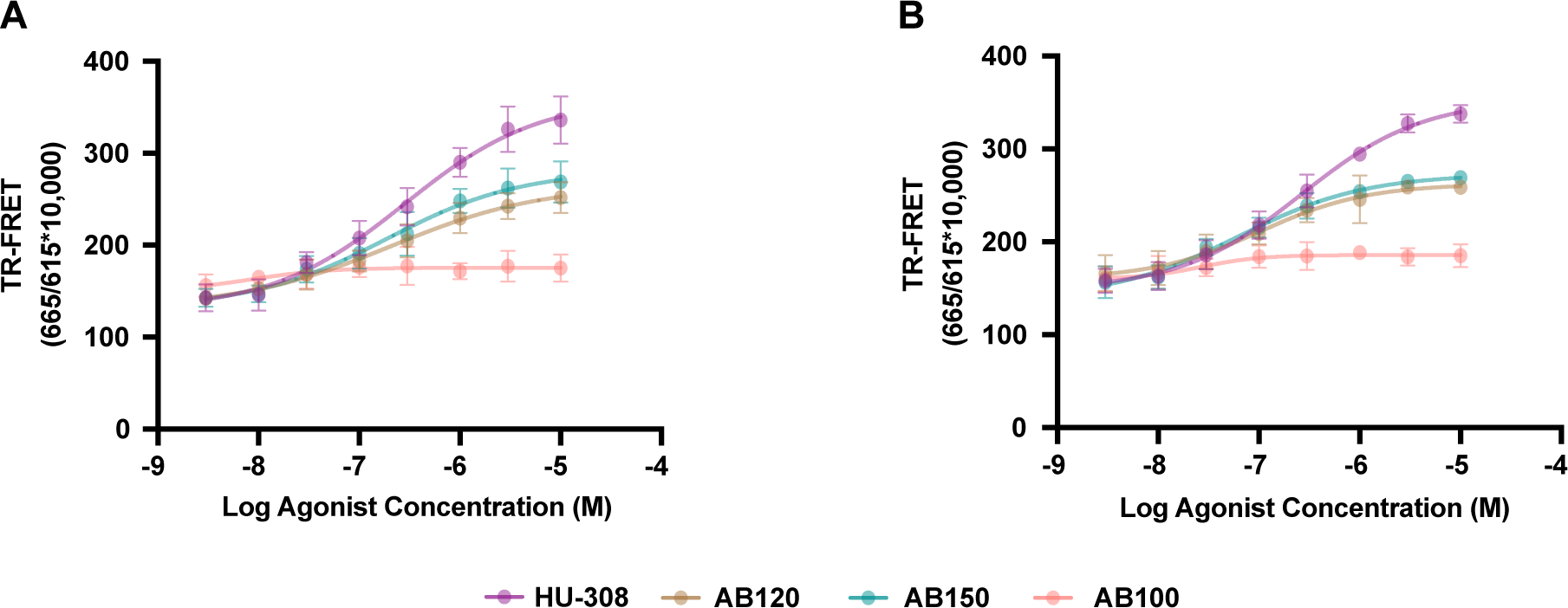
CB2 agonist antibodies-promoted β-Arrestin-2 recruitment detected by the TR-FRET assay. Dose response curve of β-Arrestin-2 recruitment by CB2 agonist antibodies in RAW264.7 (A) and THP-1 derived cells (B). Cells were transfected with 20ng of human β-Arrestin-2 expressing plasmid and 130ng empty pcDNA3 plasmid followed by measurement of β-Arrestin-2 recruitment assay. Data are represented as mean ± SD from two independent experiments.

CB2 agonists are also known to signal through a third pathway by inducing phosphorylation of ERK1/2 through G_βγ._^40^ Therefore, we measured the amount of ERK1/2 phosphorylation induced by CB2 agonists using a TR-FRET based assay. The potencies (EC_50_) of both AB120 and AB150 are similar to HU308 in both RAW264.7 and differentiated THP-1 cells and to that of the β-arrestin-2 recruitment (Figure 3 and Table 1). However, in contrast to β-arrestin-2 recruitment, both antibodies have a similar E_max_ to HU-308.

**Fig.3.**
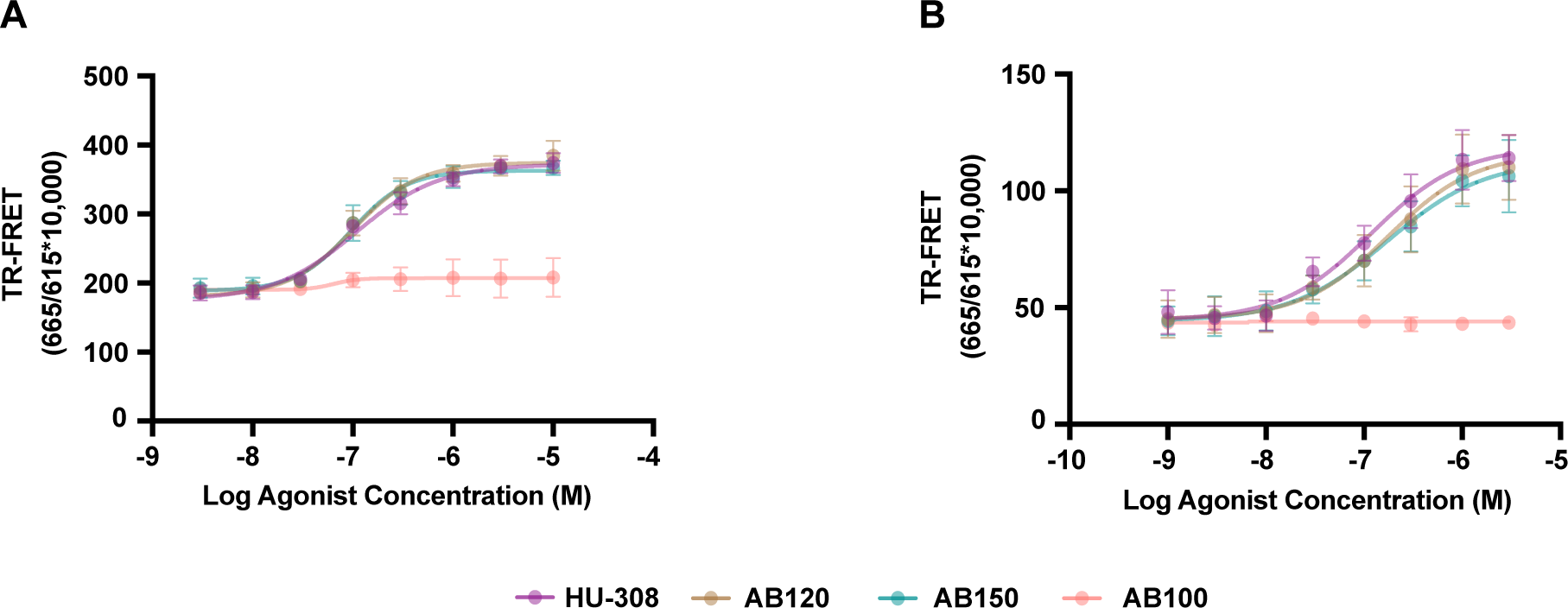
CB2 agonist antibodies-promoted ERK1/2 phosphorylation detected by the *LANCE* Phospho-ERK1/2 TR-FRET assay. (A and B) show dose dependent ERK1/2 phosphorylation by CB2 agonist antibodies in RAW264.7 and THP-1 derived cells. Data are represented as mean ± SD from two independent experiments.

Reduction of cAMP via CB2 agonism is known to prevent the activation of PKA, thereby reducing the expression of pro-inflammatory genes.^41,42^ Therefore, we next sought to study the downstream consequences of the initial signaling events in macrophages. Macrophages are a primary component of the innate host defense immune response and are involved in tissue repair and homeostasis. Depending on external stimuli, macrophages can be induced to express markers that are traditionally characterized as M1, pro-inflammatory, or M2, anti-inflammatory states.^19,23,26,43^ The M1/M2 paradigm is currently recognized as an oversimplification of the actual variation in activated macrophages; however, here it serves as a useful approach to distinguish functional effects of CB2 agonism in isolated macrophage cell lines. Considering this, an ideal CB2 agonist antibody would reduce expression of pro-inflammatory cytokines and chemokines from M1 macrophages. M2 macrophages are more complicated as they are primarily considered to be anti-inflammatory, however, upon chronic activation, M2 macrophages secrete TGF-β, which is profibrotic.^44–47^ Therefore, an ideal CB2 agonist may be one that inhibits activation of both M1 and M2 macrophages or one that biases the polarization toward M2 over M1 in the context of macrophage cell lines.

Here we demonstrate that CB2 agonism by AB120 and AB150 affects mRNA expression in mouse and human macrophage cell lines that have been stimulated with LPS or IL-4 to induce M1 or M2-like states, respectively. For RAW264.7 or differentiated THP-1 cells stimulated with LPS, CB2 agonist antibodies reduce the expression of M1 macrophage associated, pro-inflammatory genes (*TNF-α, NOS2, CCL4, IL-6,* and *IL-1β*) compared to untreated LPS stimulated and unstimulated cells (Figure 4, Table 2). Treatment of these same cell lines with IL-4 to induce M2 polarization, increases the expression of M2 macrophage associated genes (*Arg1, Mrc2, Mgl1,* and *Clec7A*) (Figure 4, Table 2). CB2 agonists HU308, AB120, and AB150 prevent the upregulation of these genes in RAW264.7 cells. However, in differentiated THP-1 cells, while HU308 and AB120 prevented the IL4 induced expression of *Arg1, Mrc2, Mgl1,* and *Clec7A,* AB150 only prevented the upregulation of *Arg1* and not *Mrc2, Mgl1,* or *Clec7A*. AB100, the isotype control, had no effect on induced M1 or M2 macrophage gene expression. Overall, these CB2 agonists influenced the M1 and M2 macrophage gene expression biasing it toward an anti-inflammatory and anti-fibrotic state with more M2 associated gene expression than M1.

**Fig.4.**
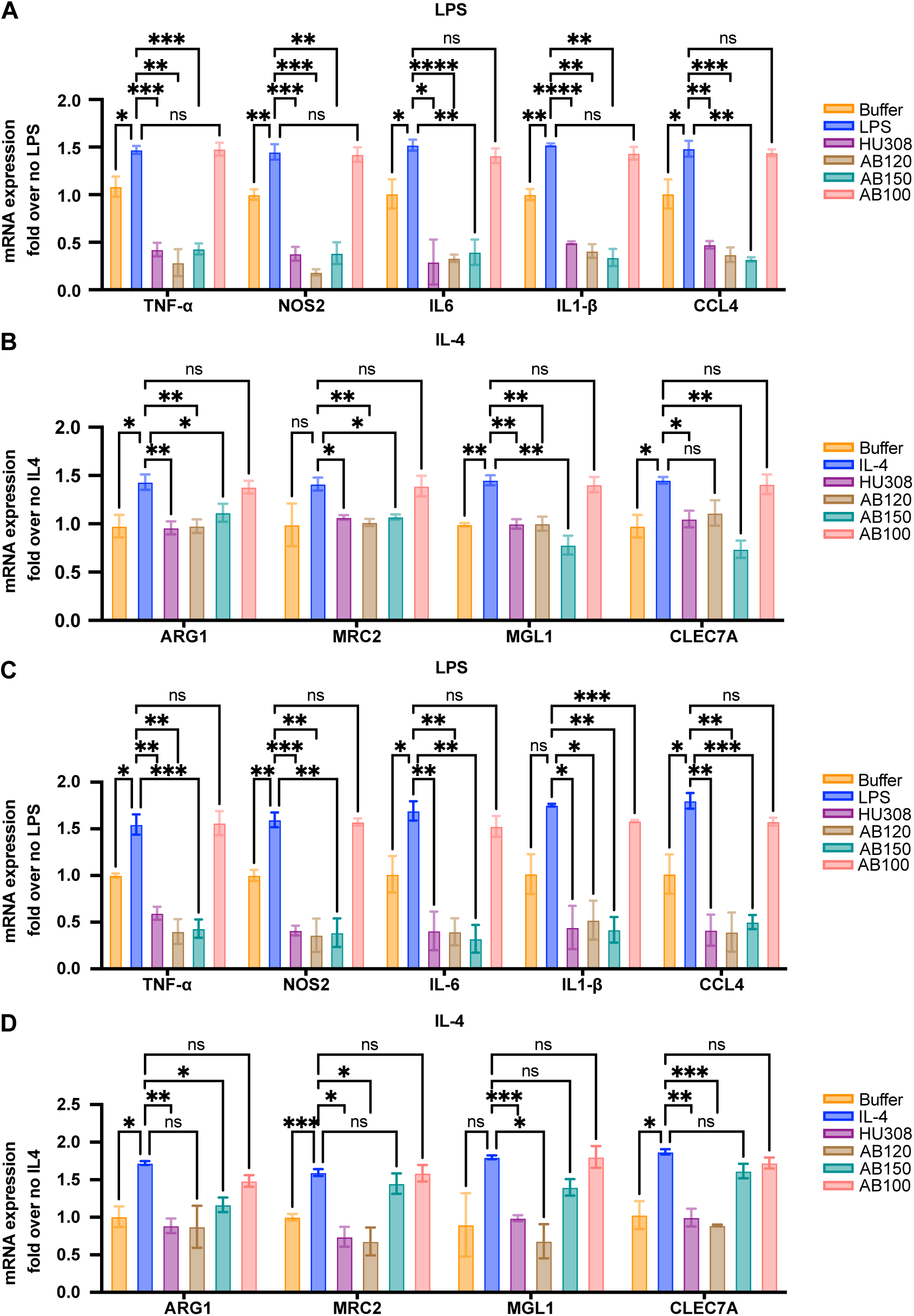
mRNA expression of M1 and M2 macrophage genes in RAW264.7 and THP-1 derived cells. Cells were incubated with LPS (100ng/ml) or IL-4 (5ng/ml). Data are represented as mean ± SD from two independent experiments. Statistical significance is defined as: * P < 0.05, ** P < 0.01, *** P < 0.001 and **** P < 0.0001

**Table 2.** Protein markers of M1 and M2 macrophage activation and their function.

| **Protein (Gene)** | **Role** | **References** |
| --- | --- | --- |
| α-SMA (*Acta2*) | Fibrogenesis, marker of HSC activation | 48 |
| ARG1 | Wound healing, marker of M2 macrophage activation | 49,50 |
| CCL4 | Chemokine, recruits immune cells to sites of inflammation | 51 |
| Dectin-1<br>(*Clec7A*) | Regulates phagocytosis, respiratory burst and autophagy | 52 |
| Collagen | Primary component of extracellular matrix and fibrosis | 53 |
| IL-10 | Anti-inflammatory cytokine | 54 |
| IL-1β | Pro-inflammatory cytokine | 55 |
| IL-6 | Pro-inflammatory cytokine | 56 |
| MGL1 | Pathogen recognition, regulate immune homeostasis | 57,58 |
| MRC2 | Collagen uptake in macrophages | 59 |
| NOS2 | Production of reactive oxygen species | 49 |
| TNF-α | Pro-inflammatory cytokine | 60 |

CB2 agonists have the potential to treat fibrosis by limiting macrophage activation, thereby reducing expression of pro-inflammatory and pro-fibrotic genes.^22,61,62^ To demonstrate the potential of AB120 and AB150 to reduce the expression of fibrotic markers, human precision cut liver slices (hPCLS) from de-identified resection samples of human liver were treated with AB120, AB150, AB100, or Alk5i (a TGF-βRI kinase inhibitor and a positive control for this assay). This ex vivo fibrosis model preserves all cell types and tissue architecture, enabling a more comprehensive study of endogenous cell-type interactions in the liver.^63,64^ Forty-eight (48) hours after treatment, mRNA expression of *Col1A1, Acta2, IL6, IL10,* and *TNF-α* and collagen secretion were measured via rtPCR and ELISA. Interestingly, both AB120 and AB150 reduced the gene expression of these fibrosis and inflammation markers, as well as the secretion of collagen as measured by ELISA (Figure 5, Table 2). As *Col1A1* and *Acta2* are markers of hepatic stellate cell (HSC) activation and collagen is primarily expressed from these stellate cells, these data demonstrate that, while AB120 and AB150 directly affect macrophage activation, they also have downstream effects on other cell types that contribute to the fibrotic process.

**Fig.5.**
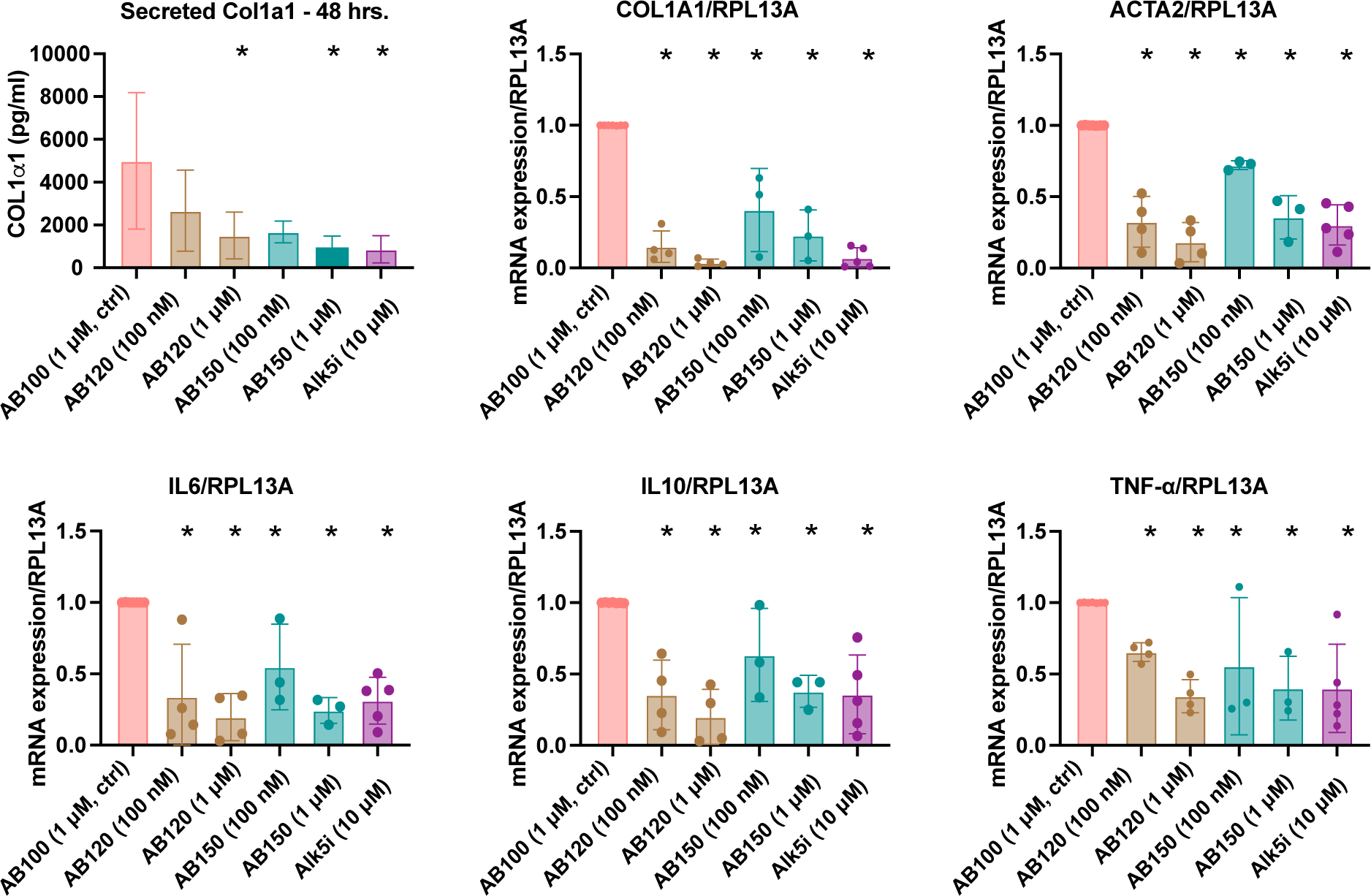
CB2 antibodies reduced expression of fibrogenic and inflammatory genes (B-F) and secretion of collagen (A) in human precision cut liver slices. For B-F, each point is the mean of mRNA expression normalized to RPL13A expression for an individual patient sample (n=3-6) measured in triplicate via rtPCR, with the bar representing the overall mean ± SD. For A, collagen in the culture medium was quantified via ELISA and depicted as mean ± SD of a single measurement per patient sample (n=3-6). Statistical significance is defined as: * P < 0.05

## Discussion

While CB2 agonism shows therapeutic potential in a number of diseases, most small molecule CB2 agonists are highly lipophilic, have short in vivo half-lives, and have unwanted activity on CB1.^8^ To address this, we have developed the first two CB2 agonist VHH-Fc antibodies, AB120 and AB150, which were discovered using a function-based selection.^10^ Antibodies that are GPCR agonists are extremely rare due to limitations of traditional antibody discovery techniques that optimize for and produce high affinity binders, which usually do not affect target function beyond inhibition. In fact, to date only seven GPCR agonist antibodies have been previously reported in the literature.^65–72^ Antibody drugs have excellent safety and efficacy track records in management of chronic conditions (e.g., the top-selling drug adalimumab [Humira] and many others^73^). Antibodies are often more target specific, with fewer off-target effects and have longer half-lives and duration of action than small molecule or peptide-based drugs. VHH domains have been clinically validated (i.e. Ablynx’s Cablivi) and their small paratope size has been hypothesized to be more compatible with GPCR epitope size than the VH/VL paratope of conventional monoclonal antibodies.^74^ Antibodies also have the advantage of peripheral restriction, which lowers potential CNS effects compared to small molecule agonists, particularly considering the CNS-biased expression of CB1 and the psychotropic effects of activation.^6^ AB120 and AB150 are VHH-Fc antibodies, which consists of the VHH linked to a modified human IgG1 Fc with reduced effector function. This VHH fusion to the Fc extends the antibody half-life in vivo.

Importantly, we have identified molecules with desired biases of downstream signaling. GPCRs, including CB2, are known to engage multiple pathways upon activation.^8,75,76^ The degree of pathway activation, or biased signaling, has downstream biological effects, which can determine the ultimate pharmacology of an agonist. Here we demonstrate cAMP biased CB2 agonism by AB120 and AB150 on mouse and human macrophage cell lines. These antibodies potently decrease NKH477 stimulated cAMP production similarly to HU-308, a CB2 selective small molecule agonist. This decrease in cAMP production ultimately reduces expression of pro-inflammatory genes by CREB phosphorylation and inhibiting NF-κB induced transcription.^42,77^ In contrast, AB120 and AB150 recruit β-arrestin-2 with lower potency and efficacy. As β-arrestin-2 recruitment can lead to internalization and down-regulation of CB2,^7,39^ this signaling bias is advantageous for a CB2 agonist therapeutic. By signaling through the G_i_/G_o_-protein pathway, with less engagement of β-arrestin, more CB2 will remain on the cell surface, enabling continued action of AB120 and AB150.

Previous reports also demonstrate that stimulation of CB2 leads to ERK-mediated cellular activation and anti-inflammatory effects in monocytes/macrophages.^40,78,79^ In agreement with these reports, we showed that AB120 and AB150 antibodies induce ERK1/2 phosphorylation in mouse and human macrophage cells suggesting CB2 agonist antibodies engage with CB2 and are coupled to the activation of the ERK-1/2 kinase cascade through a classical upstream mechanism involving G_i_/G_o_-protein βγ−subunits.^40^

Also important is the specificity of CB2 agonist therapeutics. Given that many small molecule CB2 agonists have residual CB1 activity, specificity of AB120 and AB150 for CB2 is important to avoid CB1 induced off-target pro-inflammatory and CNS effects. We demonstrate this specificity as the inhibition of cAMP production by AB120 and AB150 is counteracted by a CB2 antagonist, but not by a CB1 antagonist, in macrophage cell lines that endogenously express both CB2 and CB1.

CB2 receptors are predominantly distributed in the peripheral immune system, including macrophages, with little expression in the CNS, which makes this receptor an attractive therapeutic target for inflammation and fibrosis. Macrophages can both promote and resolve inflammation depending on their activation state and environment. In the liver, the macrophage population is heterogeneous including tissue resident macrophages (i.e. Kupffer cells) and monocyte derived macrophages.^80^ Under normal conditions, these cells maintain tissue homeostasis by balancing pro- and anti-inflammatory responses of macrophages to promote immunity to viruses, bacteria and parasites while maintaining healthy tissue through phagocytosis. Upon chronic assault with toxins, viruses, or high-fat diet, macrophages respond by polarizing to an M1 macrophage state and secreting pro-inflammatory cytokines and chemokines (CCL4, TNF-α, IL-6, IL-1β) that attract additional immune cells (i.e. neutrophils, monocytes, and NK cells), and produce reactive oxygen species (NOS2). Within the liver, these cytokines also activate hepatic stellate cells (HSC), leading to enhanced extracellular matrix (ECM) protein deposition including collagen, resulting in increased fibrosis. IL-10 is an anti-inflammatory cytokine that can counteract the effects of pro-inflammatory cytokines, however high IL-10 concentration is indicative of excess inflammation and has been linked to severity of liver cirrhosis.^81,82^ Chronic inflammation drives fibrosis that does not resolve, affecting organ function.^81^

In metabolic dysfunction-associated fatty liver disease (MAFLD), there is dysbiosis in the gastrointestinal tract that leads to increased gut permeability and higher levels of endotoxin/LPS in the serum of patients with MAFLD.^83^ LPS has been shown to stimulate the Kupffer cell (KC) secretion of multiple cytokines and chemokines, including TNF-α, IL-1β, IL-6 and the chemokine CCL4.^84^ Increased levels of these proteins leads to fibrogenic activity from hepatic stellate cells (HSC), promoting the progression of liver disease in metabolic dysfunction-associated steatohepatitis (MASH). Conversely, reduction in the levels of these proteins, as seen with AB120 and AB150 in vitro and ex vivo, should decrease HSC activation and fibrosis in MASH.

TNF-α engages HSC to increase secretion of TIMP-1, a potent contributor to liver fibrosis.^85,86^ Moreover, activation of the IL-6 pathway in conjunction with TNF-α leads to strongly pro-inflammatory effects^87^ that can synergistically trigger chronic inflammation and fibrosis in the liver. IL-1β is another potent cytokine that is increased in LPS treated macrophages that can act on multiple cell types to promote release of chemokines such as CCL2 and CCL5 for further immune cell recruitment, and can directly activate HSC to increase fibrotic activity.^88,89^ The increase in CCL4 chemokine secretion with LPS serves as a chemoattractant for multiple immune cell populations, including natural killer cells, monocytes, T cells and dendritic cells, which respond to CCL4 gradients in response to tissue damage.^90^ Furthermore, CCR5, which is a receptor for CCL4, is expressed on HSC, and activation of this receptor can directly promote hepatic fibrosis.^91^ Together, these data demonstrate that KC secretion of these multiple cytokines and chemokines can serve as a focal point for the activation of HSC and promotion of the fibrotic phenotype.

Consistent with this mechanism, we show LPS-driven increase of TNF-α, IL-1β, IL-6 and the chemokine CCL4 in two macrophage cell lines. Importantly, we have demonstrated that CB2 agonist antibodies, AB120 and AB150, decreased the expression of these LPS-induced pro-inflammatory cytokines and chemokines below baseline levels in these mouse and human macrophage cell lines. Similar reduction in inflammatory cytokines was seen with primary human tissue in hPCLS, consistent with the proposed model where reduction in these cytokines, will also reduce hepatic stellate cell activation and subsequently reduce collagen expression and deposition. This model is further supported by the reduced expression of smooth muscle actin and alpha-1 type I collagen observed upon treatment with AB120 and AB150 in the hPCLS model.

On the other hand, treatment of macrophages with IL4 promotes expression of anti-inflammatory M2 macrophage markers. However, when chronically activated, M2 macrophages can be pro-fibrotic due to increased expression of TGF-β.^50,92^ On mouse macrophages, AB120 and AB150 prevented IL-4 induced expression of M2 macrophage markers, but did not decrease the expression from baseline. However, for the differentiated THP-1 cells, the effect of these CB2 agonist antibodies was varied, with AB120 preventing the IL-4 induced M2 marker upregulation and AB150 only preventing *Arg1* upregulation. Overall, using the simplified M1/M2 model, these antibodies bias macrophage polarization to favor anti-inflammatory over pro-inflammatory gene expression. While more complex in vivo, this biased polarization is an advantage for these antibodies in treating liver fibrosis and resolving the inflammatory state.

In conclusion, we identified the first agonist antibodies for CB2 - one of only eight GPCRs for which antibody agonists have been discovered - and in vitro studies validate our hypothesis that our first-in-class CB2 agonist antibodies signal through G-alpha biased signaling and reduce expression of pro-inflammatory cytokines and macrophage activation. The effect on in vitro signaling, gene expression and macrophage activation directly translates to gene expression in human liver tissue, where reduction in HSC activation, inflammation and fibrosis markers are also observed in the hPCLS studies. These data make a strong case for AB120 and AB150 as promising therapeutics for liver fibrosis.

In addition to liver fibrosis, macrophages drive other fibrotic disease including pulmonary and renal fibrosis.^22^ CB2 is also expressed on peripheral neurons and microglial cells; therefore, these CB2 agonists have potential applications for treating peripheral neuropathies induced by chemotherapy or diabetes.^93–95^ Based on the functional biology demonstrated by AB120 and AB150 in this report, we are pursuing further evaluation of these antibodies for therapeutic use in multiple indications, including liver fibrosis and peripheral neuropathies.

## Materials and Methods

### Cell lines, reagents and ligands

Mouse macrophage cell line (RAW264.7) and Human leukemia monocytic cell line (THP-1) cell lines were purchased from the American Type Culture Collection (ATCC) and cultured in Dulbecco’s modified Eagle medium (DMEM) and RPMI-1640 medium from ATCC supplemented with 10% FBS, 20mM HEPES and 1X antibiotic and antimycotic, respectively. All cells were maintained at 37 °C in the presence of 5% CO_2_. 3-Isobutyl-1-methylxanthine, LPS and IL-4 are from Sigma-Aldrich. RO-1276, NKH 477, HU-308, SR 155528, ACEA, SR141716A (rimonabant), and Phorbol 12-myristate 13-acetate (PMA) are from Tocris.

### Antibody cloning and production

VHH nucleotide sequences for AB100, AB120, and AB150 were procured as gene fragments from IDT and cloned into the TGEX-Fc expression vector (Antibody Design Labs) using NEBuilder HiFi DNA assembly (New England Biolabs). For each antibody, plasmid DNA was transfected into Expi 293F cells (ThermoFisher Scientific) with Expi 293 transfection reagent using the manufacturer’s protocol and incubated at 37^0^C with 80% CO2 for 5 days. The protein A resin, MabSelect Sure (Cytiva), was used to purify the VHH-Fc fusions from the cell culture supernatant through gravity flow. The purified antibodies were dialyzed into 1X PBS (pH 7.4).

### Differentiation of THP-1 and primary human monocyte-derived macrophages

THP-1 cells were seeded at 0.25×10^6^ cells/mL and stimulated with 200ng/mL phorbol 12-myristate 13-acetate (PMA, Tocris) for 24 hrs. Then, culture media was removed, and fresh media was added. After 96 hrs, THP-1 cells are fully matured into macrophages and firmly attached to the bottom of the culture vessel (Supplementary Figure 1). Cells can be now used for subsequent experiments.^96^

### Microscope image acquisition

THP-1 cells were seeded on 6 well plate and differentiated with PMA. A Nikon Eclipse TE300 Inverted Microscope equipped with a Hg short arc HBO lamp (OSRAM) was used to acquire images. The phase-contrast images were collected using a 10x objective lens. The images were collected before and five days after PMA treatment.

### Luminescence cAMP assay

cAMP-Glo Max assay kit (Promega, V1682) was used to measure cAMP signals in Raw264.7 and THP-1 cells. Briefly, 10000 cells per well seeded on Poly-D-lysine (Sigma Aldrich) coated 384 well view plate (Revvity) and incubate 16-18 hours. Test compounds were added to the wells, briefly centrifuged, and incubated for 30 min at 37°C. After 30 min, cAMP detection reagent (Glo one) with PKA was added to cells, briefly centrifuged, and incubated 20 min at ambient temperature (RT). In the final step, kinase Glo reagent was added to the cells, briefly centrifuged, and incubated for 10 min at RT. The luminescence signal was measured by using Synergy HTX multi-mode microplate reader (Bio Tek). The data were fit with a non-linear regression of variable slope using GraphPad Prism software (version 10.0)

### TR-FRET Phospho-ERK Detection assay

The TR-FRET Phospho-ERK assay was carried using LANCE Ultra phospho-ERK1/2 (Thr202/TYR204) detection kit (Revvity, TRF400C) following manufacturer’s protocol. Briefly, 80,000 cells were seeded on 96 well clear bottom plates and incubated overnight at 37°C. The next day, cells were serum starved for 4 hrs. followed by removal of the serum free medium. Test compounds were then added to the plate and incubated 5 min at RT with shaking (200rpm). Later, test compounds were removed and lysis buffer added to the plate and incubated for 30 minutes at RT with shaking (200 rpm). Cell lysates (15ul) were transferred to 384 Optiview detection plates (Revvity) and premixed Eu-labeled anti-ERK1/2 (T202-Y204) antibodies and ULight labeled anti-ERK1/2 antibodies were added to the plate and incubated 1hr at RT. TR-FRET signal was measured with a Victor2 multi plate reader. The ratio ((em.655/em615)*10,000) was calculated and the data fit with a non-linear regression of variable slope using GraphPad Prism software (version 10.0).

### β-arrestin-2 recruitment assay

The β-arrestin-2 recruitment assay was performed using HTRF β-arrestin 2 Recruitment Detection Kit (Revvity) following manufacturer’s protocol. Briefly, cells were seeded on 96 well culture plate and incubated for 24 hrs at 37 °C. After 24hrs, β-arrestin (20ng) and empty pcDNA3 (150ng) plasmids were transfected to the cells by using lipofectamine 2000 transfection kit (Thermo Fisher Scientific) and incubated 24hr at 37°C incubator. After 24hrs, the media was removed, and test compounds were added to the plate and incubated for 30min. Later, the media was removed, and stabilization buffer was added to the plate and incubate for 15 min at RT. Stabilization buffer was removed, and the plate was washed 3x with wash buffer. Later, pre-mixed d2 and Eu cryptate antibodies were added to the cells and incubated overnight at RT. Next day, HTRF signal was measured by using a Victor2 multi plate reader. The ratio ((em.655/em615)*10,000) was calculated and fit the data with a non-linear regression of variable slope using GraphPad Prism software (version 10.0).

### Quantitative PCR

Macrophage gene expression quantification was adapted from previous publications.^19,23^ Cells were seeded on 6 well culture plate and incubate 24hrs at 37°C incubator. Next, cells were serum starved for 24hrs and added test compounds to the cells and incubated 16 to 18hr at 37°C. After 18hrs, LPS (100ng/ml) or IL-4 (10ng/ml) was added to the cells, followed by 8hrs and 16hrs incubation, respectively. After incubation, cells were lysed and RNA extracted from collected samples using RNeasy Mini Kit (Qiagen) according to the manufacturer’s protocol. mRNA was reverse transcribed using Super script IV VILO master mix kit (Invitrogen). Finally, quantitative PCR (qPCR) was performed on AriaMx RT-PCR system (Agilent), using Power SYBR Green Universal Master mix (Applied Biosystems). Samples were measured in triplicates. The qRT-PCR data were processed and analyzed by the 2^-ΔΔCT^ method. The sequences of the primer pairs used are listed in Supplementary Table 1.^19^ Statistical analyses were performed with GraphPad Prism software (version 10.0) by using two-way ANOVA with Dunnett’s multiple comparisons test. Statistical significance is defined as: * P < 0.05, ** P < 0.01, *** P < 0.001 and **** P < 0.0001

### Precision cut liver slices

The technique for human precision-cut liver slices (hPCLS) was previously described,^64^ with minimal modification. The protocol relies on fully de-identified liver samples and thus formal review by the Institutional Review Board (IRB) at the School of Medicine at Mount Sinai is not required. Briefly, de-identified patient liver samples surrounding HCC or metastatic regions were transported to the lab in Belzer UW Cold Storage Solution (UW Solution). An 8 mm diameter core was obtained, and 200 μm slices were generated from each core in the presence of UW solution, with continuous supplies of 95% O2 and 5% CO2. The slices were cultured in complete media containing WE-GlutaMAX, supplemented with 25 mM glucose, 15% Cellartis Hepatocyte Maintenance Media (Takara Bio), 1X insulin-transferrin-selenium, 100 nM dexamethasone, 2% FBS, and 50 μg/ml gentamycin. After 24 hours of initial culture, the media were replaced with fresh complete media containing either IgG or two different doses of antibodies for an additional 48 hours. Conditioned media was collected for ELISA measurement of secreted Col1a1. Slices were also collected for total RNA extraction and measurement of mRNA expression. Data was normalized to treatment with isotype control (AB100). Statistical analyses were performed with GraphPad Prism software (version 10.0) by using two-way ANOVA with Dunnett’s multiple comparisons test for mRNA expression and ordinary one-way ANOVA with Dunnett’s multiple comparisons test for secreted Col1a1. Statistical significance is defined as: * P < 0.05

## Abbreviations

CB1: Cannabinoid Receptor 1,
CB2: Cannabinoid Receptor 2,
CNS: Central nervous system,
CREB: cAMP-response element-binding protein,
ECM: Extracellular matrix,
ERK: Extracellular signal-regulated kinase,
GPCR: G-protein coupled receptor,
HSC: Hepatic stellate cells,
hPCLS: human precision cut liver slices,
LPS: Lipopolysaccharide,
MAFLD: Metabolic dysfunction-associated fatty liver disease,
MASH: Metabolic dysfunction-associated steatohepatitis,
PKA: Protein kinase A,
TGFβ: Transforming growth factor beta,
TR-FRET: Time-Resolved Fluorescence Resonance Energy Transfer

---

**Supplementary Figure1.**
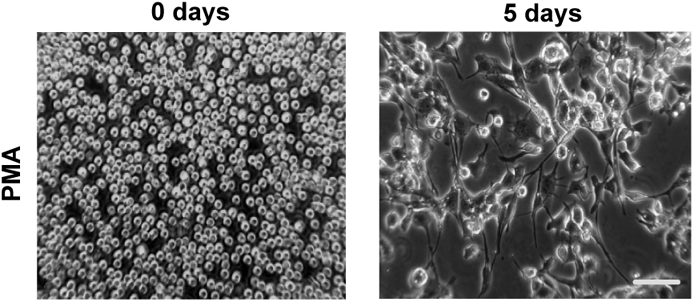
THP-1 Cells differentiation by PMA. THP-1 cells were incubated for 24h without (A) or with 200ng/mL PMA (B) and replaced the media. Images were captured at 10x magnification after 5 days by using phase-contrast inverted microscope. Scale bar −10μm

## Data Availability

The authors declare that all the data supporting the findings of this study are available within the paper and its Supplemental Data.

## Acknowledgements

We thank Rodrigo Baltanas, Brett Robinson, Gustavo Pesce, Sameer Soi, Stephanie Lopez and Allison Cooke for their work to develop Abalone Bio’s antibody discovery platform and to discover CB2 agonist antibodies. We also thank Ken Mackie and Amey Dhopeshwarkar at Indiana University for their support in functional assessment of CB2 agonist antibodies. Finally, we thank Alejandro Colman-Lerner for his continued support and yeast genetics expertise.

## Author Contributions

Raghavender Reddy Gopireddy – Investigation, methodology, formal analysis, visualization, writing – original draft, writing-review and editing

Dipankar Bhattacharya – Investigation, formal analysis

Swastik Sen – Investigation, resources, writing – original draft

Monica Schwartz – Investigation, supervision, writing-review and editing

Scott L. Friedman – Methodology, funding acquisition, writing-review and editing

Richard Yu – Conceptualization, funding acquisition, methodology, writing-review and editing

Toshi Takeuchi – Supervision, methodology, writing-review and editing

Lauren Schwimmer – Supervision, formal analysis, methodology, funding acquisition, writing – original draft, writing – review and editing

## Declaration of generative AI and AI-assisted technologies in the writing process

During the preparation of this work the authors used Gemini 2.5, ChatGTP Teams, and Claude Sonnet 4 to review the readability and language of the text. After using these tools, the authors edited the text as needed and take full responsibility for the content of the published article. The text was not directly edited by these AI technologies.

---

**Supplemental Table 1.**
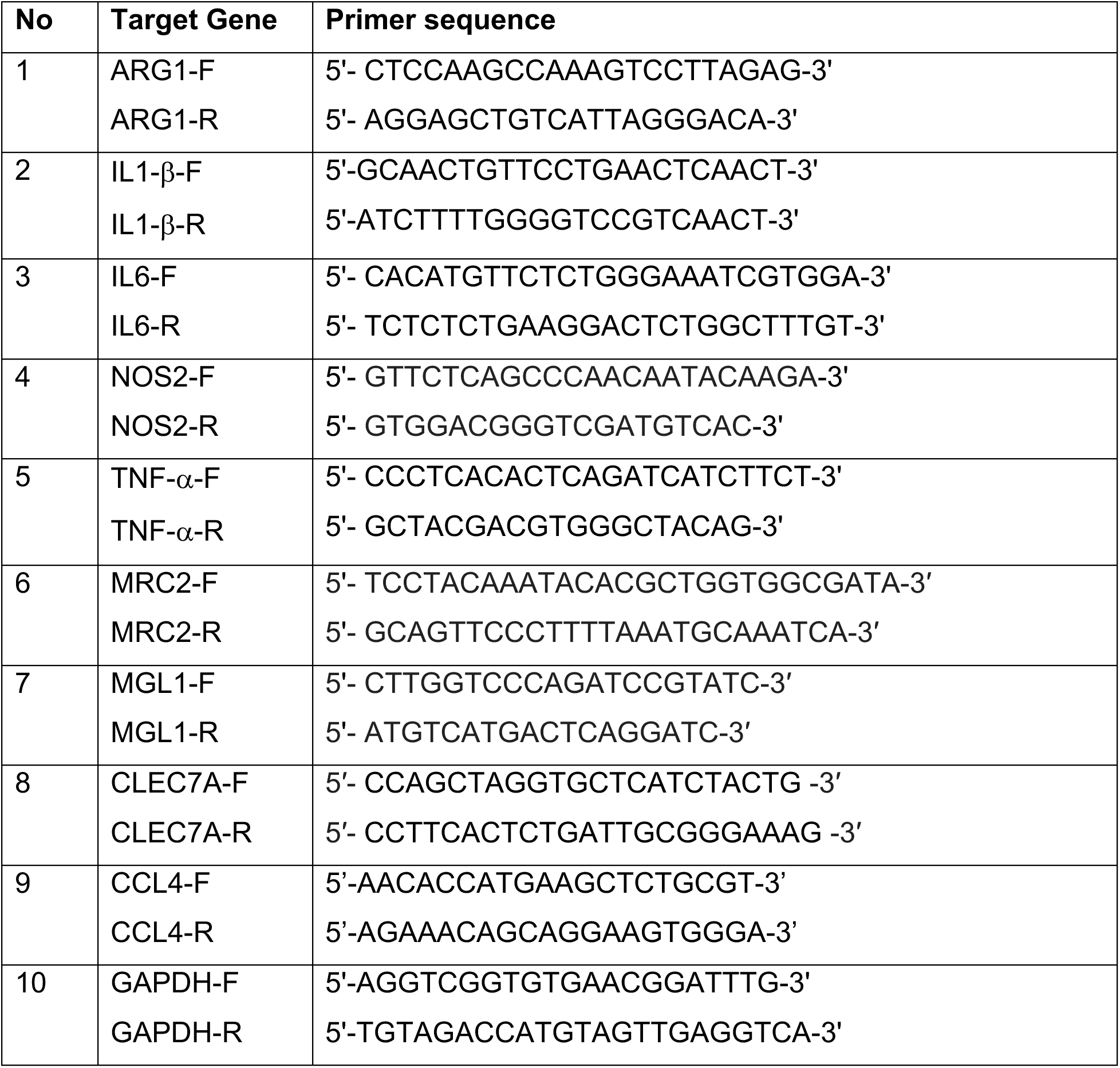

